# Molecular mechanisms for intestinal HCO_3_^-^ secretion and its regulation by guanylin in seawater-acclimated eels

**DOI:** 10.1101/580761

**Authors:** Yoshio Takei, Marty K.S. Wong, Masaaki Ando

## Abstract

The intestine of marine teleosts secretes HCO_3_^-^ into the lumen and precipitates Ca^2+^ and Mg^2+^ in the imbibed seawater as carbonates to decrease luminal fluid osmolality and facilitate water absorption. However, reports on studies on the hormonal regulation of HCO_3_^-^ secretion are just emerging. Here, we showed that guanylin (GN) applied to the mucosal side of intestinal epithelia increased HCO_3_^-^ secretion in seawater-acclimated eels. The effect of GN on HCO_3_^-^ secretion was slower than that on the short-circuit current, and the time-course of the GN effect was similar to that of bumetanide. Mucosal bumetanide and serosal 4,4’-dinitrostilbene-2,2’-disulfonic acid (DNDS) inhibited the GN effect, suggesting an involvement of apical Na^+^-K^+^-2Cl^-^ cotransporter (NKCC2) and basolateral Cl^-^/HCO_3_^-^ exchanger (AE)/Na^+^-HCO_3_^-^ cotransporter (NBC) in the GN effect. However, mucosal DNDS and diphenylamine-2-carboxylic acid (DPC) failed to inhibit the GN effect, showing that apical AE and Cl^-^ channel are not involved. To identify molecular species of possible transporters involved in the GN effect, we performed RNA-seq analyses followed by quantitative real-time PCR after transfer of eels to seawater. Among the genes upregulated after seawater transfer, those of Slc26a3a, b (DRAa, b) and Slc26a6a, c (Pat-1a, c) on the apical membrane of the intestinal epithelial cells, and those of Sls4a4a (NBCe1a), Slc4a7 (NBCn1), Slc4a10a (NBCn2a) and Slc26a1 (Sat-1) on the basolateral membrane were candidate transporters involved in HCO_3_^-^ secretion. Judging from the slow effect of GN, we suggest that GN inhibits NKCC2b on the apical membrane and decreases cytosolic Cl^-^ and Na^+^, which then activates apical DNDS-insensitive DRAa, b and basolateral DNDS-sensitive NBCela, n1, n2a to enhance transcellular HCO_3_^-^ flux across the intestinal epithelia of seawater-acclimated eels.

## Introduction

Marine teleosts drink seawater (SW) copiously and absorb >80% of imbibed SW across the intestine to compensate for osmotic water loss in the hypertonic environment (Grosell, 2011). The ingested SW is diluted to isotonicity by coordinated actions of NaCl removal by the esophagus (Hirano and Meyer-Gostan, 1976; Parmelee and Renfro, 1983; Takei et al., 2017) and Ca^2+^/Mg^2+^ removal by precipitation as carbonate (Grosell et al., 2009b; Wilson et al., 2009; Faggio et al., 2011). Water is then absorbed by the intestine in parallel with NaCl absorption from diluted SW (Grosell, 2011). The molecular mechanisms for NaCl absorption across the intestinal epithelia have been well characterized in a few teleost species; Na^+^-K^+^-Cl^-^ cotransporter 2 (NKCC2 or Slc12a1) is the major player for NaCl uptake on the apical membrane of enterocytes and Na^+^/K^+^-ATPase (NKA) and unidentified Cl^-^ channel on the basolateral membrane transport NaCl into the extracellular fluid (Cutler and Cramb, 2008; Gregŏrio et al., 2013; Ando et al., 2014; Esbaugh and Cutler, 2016). The molecular mechanism for HCO_3_^-^ secretion has also been investigated in the intestine of marine teleosts (Grosell, 2011). The major transporter for HCO_3_^-^ secretion on the apical membrane was identified as Pat-1 (Putative anion transporter-1)/Slc26a6 and that for HCO_3_^-^ uptake on the basolateral side was suggested as electrogenic Na^+^- HCO_3_^-^ cotransporter (NBCe1/Slc4a4) because of upregulation of these genes after high salinity acclimation in the pufferfish (Kurita et al., 2008), toadfish (Grosell et al., 2009b; Taylor et al., 2010) and seabream (Gregŏrio et al., 2013). The combination of these transporters accomplishes transcellular HCO_3_^-^ secretion.

Intestinal ion and water transport is regulated by various hormones (Takei and Loretz, 2011), of which intestinal guanylin (GN) family peptides are promising candidates for the major hormone involved in SW adaptation (Kalujnaia et al., 2009; Takei and Yuge, 2007). We have identified GN, uroguanylin (UGN) and renoguanylin (RGN), and two guanylyl cyclase-C receptors (GC-C1 and GC-C2) in the intestine of Japanese eel (Yuge et al., 2003, 2006). These hormone and receptor genes are expressed abundantly in the different segments of digestive tract, and importantly, their expression is consistently upregulated after transfer of fish from fresh water (FW) to SW. Among the guanylin family, only GN is intestine-specific and secreted from goblet cells into the lumen with mucus (Yuge et al., 2003). Because homologous GN from the eel is available, this euryhaline teleost serves as a good model to analyze the mechanisms of GN action on the HCO_3_^-^ secretion. It was shown that GN decreased NaCl absorption through inhibition of NKCC2b (Ando et al., 2014) and decreased Cl^-^ secretion into the lumen through stimulation of DPC-sensitive anion channels in the intestine of SW-acclimated eels (Ando and Takei, 2015). In the marine toadfish, eel RGN was shown to decrease HCO_3_^-^ secretion through inhibition of Slc26a6 in the posterior intestine, while it increased the secretion in the fish acclimated to concentrated (60 ppt) SW (Ruhr et al., 2015). In view of the highly diversified osmoregulatory mechanism and their hormonal regulation in fishes (Takei et al., 2014), it is desirable to examine how homologous GN affects the suite of transporters for HCO_3_^-^ secretion in the eel.

In this study, we examined the effect of GN on HCO_3_^-^ secretion in the intestine of SW-acclimated eels using pH stat method in the Ussing chamber. As GN increased HCO_3_^-^ secretion, we examined the effect of various inhibitors for transporters on the GN-induced HCO_3_^-^ secretion to probe the mechanism. To identify target transporters, we then performed RNA-seq analyses and listed all candidate genes expressed in the eel intestine for HCO_3_^-^ secretion. Molecular species of the transporters were further narrowed down by quantitative real-time PCR using paralog-specific probes based on the upregulation after transfer of eels from FW to SW. Among upregulated genes, we finally selected candidate transporters involved in GN-induced transcellular HCO_3_^-^ secretion based on sensitivity to the inhibitors.

## Materials and Methods

### Animals and drugs

Cultured eels (*Anguilla japonica*) weighing around 200 g were purchased from a pond culture in Yamakine, Chiba, and kept in freshwater aquaria at 18°C for more than 1 week. Some eels were then transferred to seawater (SW) tanks and kept there for more than one week before experimentation (termed SW eels). Fish were not fed after purchase. All experiments performed in this study were approved by the Animal Experiment Committee of The University of Tokyo and performed according to the guidelines prepared by the Committee.

Eel GN (Peptide Institute, Osaka, Japan), bumetanide (Sankyo Co., Tokyo), DNDS and DIDS (Tokyo Chemical Industry Co., Ltd, Tokyo), and DPC (Wako Pure Chemical Industries Ltd., Tokyo) were purchased from commercial sources (see footnote for abbreviations). GN was dissolved in water at 10^-4^ M, aliquoted by 100 μl, kept frozen at −20°C, and diluted by the appropriate Ringer solutions at the time of experimentation. Bumetanide and DPC were dissolved in ethanol at 10^-2^ M, and final concentration of ethanol was less than 1% after dilution with HCO_3_^-^-free Ringer. One percent ethanol alone had no effects on the electrical parameters of tissues in the Ussing chamber. DNDS and DIDS were dissolved in HCO_3_^-^-free Ringer (for mucosal side addition) or in normal Ringer (for serosal side) at 10^-2^ M, and diluted to 5 x 10^-4^ M before administration.

### Physiological studies using Ussing chamber

After decapitation, the intestine of SW eels (n=31 in total) was removed and stripped of the serosal muscle layers. The stripped intestine was opened and mounted as a flat sheet in an Ussing chamber with an exposed area of 0.785 cm^2^. The serosal side of the intestine was bathed with Ringer solution (2.3 ml), containing (in mM) 118.5 NaCl, 4.7 KCl, 3.0 CaCl_2_, 1.2 MgSO_4_, 1.2 KH_2_PO_4_, 24.9 NaHCO3, 5.0 glucose and 5.0 alanine, and was bubbled with a 95% O_2_ – 5% CO_2_ gas mixture (pH 7.4). Although eel plasma pH is ca. 7.8, all experiments were performed at pH 7.4 because electrical parameters were more stable (Ando et al., 2014) and the effects of GN and bumetanide were more reproducible at this pH (Ando and Takei, 2015). The mucosal side of the intestine was bathed with unbuffered Ringer solution (2.3 ml), where NaHCO_3_ was replaced by NaCl and gassed with 100% O_2_ passed through 20 mM NaOH. The mucosal pH was measured by a pH electrode (MI-410, Microelectrodes. Inc., Bedford, USA) and a pH meter (HM-5B, Toa Electronics, Tokyo, Japan), and clamped at pH 7.4 by pH-stat (HSM-10A, Toa Electronics, Tokyo).

### HCO_3_^-^ secretion measurement

The rate of HCO_3_^-^ secretion was estimated by the rate of mucosal alkalinization, which was calculated from the amount of 50 mM HCl titrated into the chamber solution to clamp the mucosal pH. The PD was measured at the serosa relative to the mucosa through a pair of electrodes (EK1, WPI, Sarasota, USA) and a voltage/current clamp system (CEZ-9100, Nihon Kohden, Tokyo). To determine the tissue resistance (Rt), rectangular pulses (20-30 μA, 500 ms) were applied across the intestinal epithelium every 5 min. These data were recorded automatically on a chart recorder (EPR-121A, Toa Electronics, Ltd., Nagoya, Japan). From the deflection of the PD (ΔPD), total resistance was calculated and the Rt was obtained by subtracting fluid resistance (120 ohm. cm^2^) from the total resistance. The short-circuit current (Isc) was calculated as ΔPD/Rt.

### Effects of inhibitors on GN-induced HCO_3_^-^ secretion

To assess the ion transporter target(s) of GN that lead to increased HCO_3_^-^ secretion, various inhibitors were applied on either mucosal or serosal side of the intestinal epithelia in the Ussing chamber before or after GN administration. The inhibitors applied on the mucosal side were bumetanide, DPC and DNDS/DIDS since, in our earlier studies, GN inhibited NKCC2 and stimulated Cl^-^ channel in the eel (Ando et al., 2014; Ando and Takei, 2015), and since UGN inhibited Pat-1 in the toadfish (Ruhr et al., 2016). DNDS/DIDS was also applied to the serosal side to evaluate the role of basolateral AE and NBC that transport HCO_3_^-^ into the cell.

### Transcriptomic (RNA-seq) analyses

In order to identify all candidate transporters that are expressed in the eel intestine and could be involved in HCO_3_^-^ secretion, we performed RNA-seq analyses using the methods that we reported previously (Wong et al., 2014). For this purpose, eels were transferred directly from FW to SW and the intestine was collected before and 7 d after transfer (n=5 at each time point). After total RNA extraction using Isogen (Takara Bio Inc., Shiga, Japan), the RNA quality was monitored using an Agilent 2100 Bioanalyzer system (Agilent Technology, CA) and only RNA with integrity number >7.0 were used for subsequent procedures and sequencing. The cDNA libraries were prepared from the RNA samples using TruSeq RNA Sample Preparation v2 and sequenced by Illumina HiSeq 2500 (Illumina Inc., CA). The detailed protocol for bioinformatic analysis was described in Wong et al. (2014).

### Quantitative analyses of gene expression

In this experiment, the anterior, middle and posterior parts of the intestine were separately collected at 0, 3, 12, 24 h and 3, 7 d after FW-FW and FW-SW transfer (n=6 at each time point). The tissues were immediately frozen, and total RNA was extracted and treated with DNase I (Life Technology, CA) to remove genomic DNA contamination. Then, 1 μg of RNA was reverse-transcribed with Iscript cDNA Synthesis Kit (Bio-Rad, CA) according to the manufacturer’s protocols. Real-time PCR was performed using Kappa SYBR 2X PCR mix (Kappa Biosystems, MA) and ABI 7900HT Fast Real Time PCR System (Life Technology). The amplification of a single amplicon was confirmed by analyzing the melting curve after the real time cycling. Eel elongation factor 1α (*eef1a*) was used as an internal control to normalize the gene expressions among different samples. We also measured the expression of other transporter genes such as *nkcc2b* known to be upregulated in SW eel intestine to confirm the reliability of the assay. Primer sequences are listed in Supplementary Table 1.

### Statistical analyses

Statistical analyses of the data were performed using Wilcoxon signed rank test or Mann-Whitney *U*-test, programmed by KyPlot (Ver. 5.0, Kyens Lab Inc. Tokyo, Japan). Results are given as mean ± SEM and considered significant at *P* < 0.05.

## Results

### Effects of GN on HCO_3_^-^ secretion

The baseline values before treatments were −25.1 ± 3.2 mV for potential difference (PD), −501.8 ± 63.1 μA for short-circuit current (Isc), and 21.9 ± 1.4 ohm.cm^2^ for transepithelial resistance (Rt) at the middle intestine (n=13). After application of GN to the mucosal side, serosa-negative PD and short-circuit current Isc decreased within 5 min, but HCO_3_^-^ secretion increased slowly with a latency of ca. 20 min (Fig. 1A). The GN effect on HCO_3_^-^ secretion was segment-dependent with the greatest effect on the early segment of middle intestine (Fig. 1B). The GN-induced HCO_3_^-^ secretion was accompanied by a remarkable inhibition of the Isc. However, no correlation was observed between GN-induced HCO_3_^-^ secretion and Isc (Fig. 2).

**Fig. 1.**
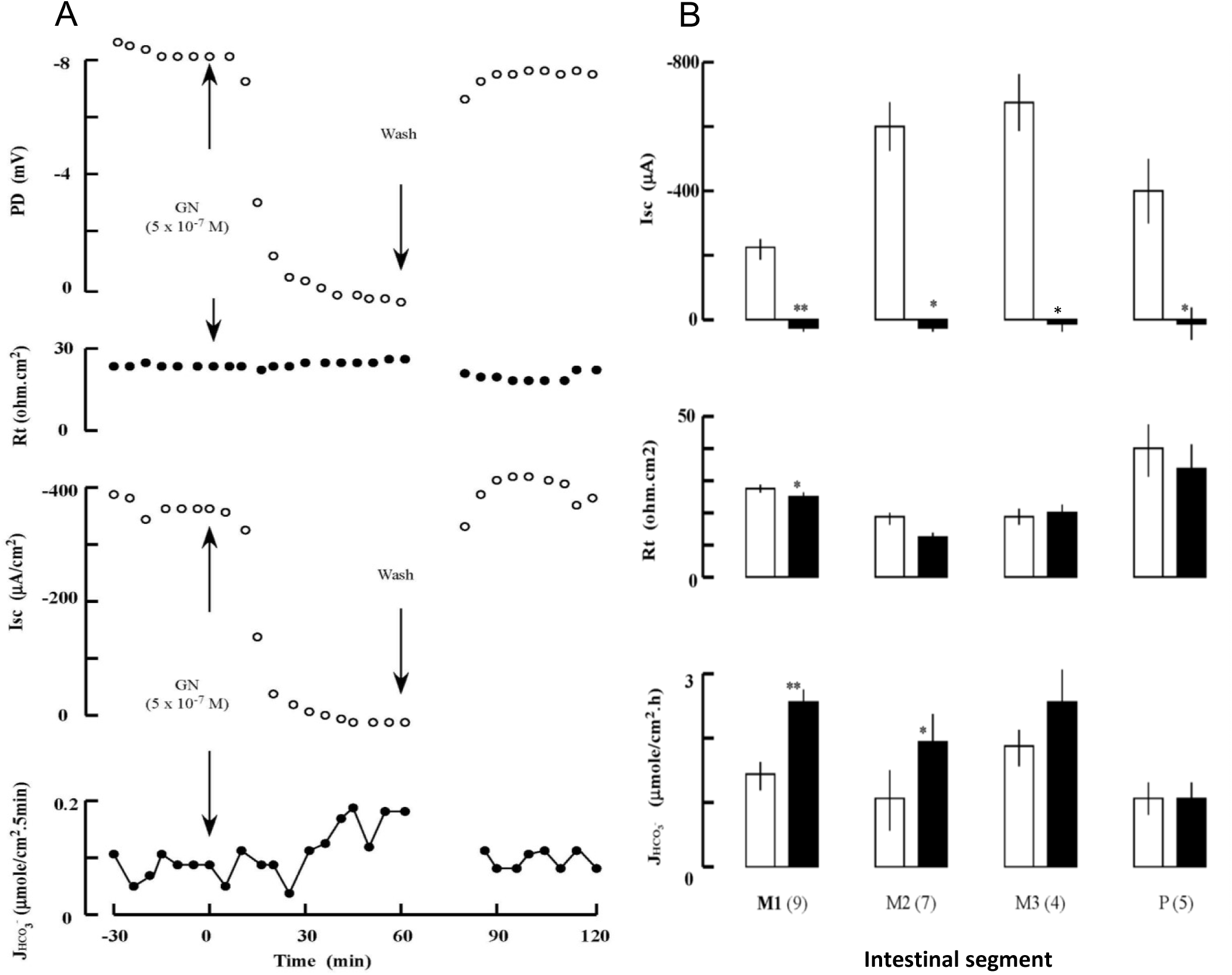
(A) An example of the effects of mucosal guanylin (GN) on the transepithelial potential difference (PD), tissue resistance (Rt), short-circuit current (Isc), and HCO_3_^-^ secretion rate JHCO_3_^-^ in the middle intestine of SW eels. Recovery from GN treatment was observed after rinsing. (B) Regional differences in the effects of GN (5 x 10^-7^ M) on Isc, Rt, and JHCO_3_^-^ in SW eel intestine. Middle and posterior segments, where GN effects were prominent, were divided as M1-3 and P as described previously (Ando et al., 2014). White and black columns show before and after treatment, respectively. Values are means ± SEM. Numbers of experiments are in parenthesis. *p<0.05, **p<0.01 compared with the value before treatment.

**Fig. 2.**
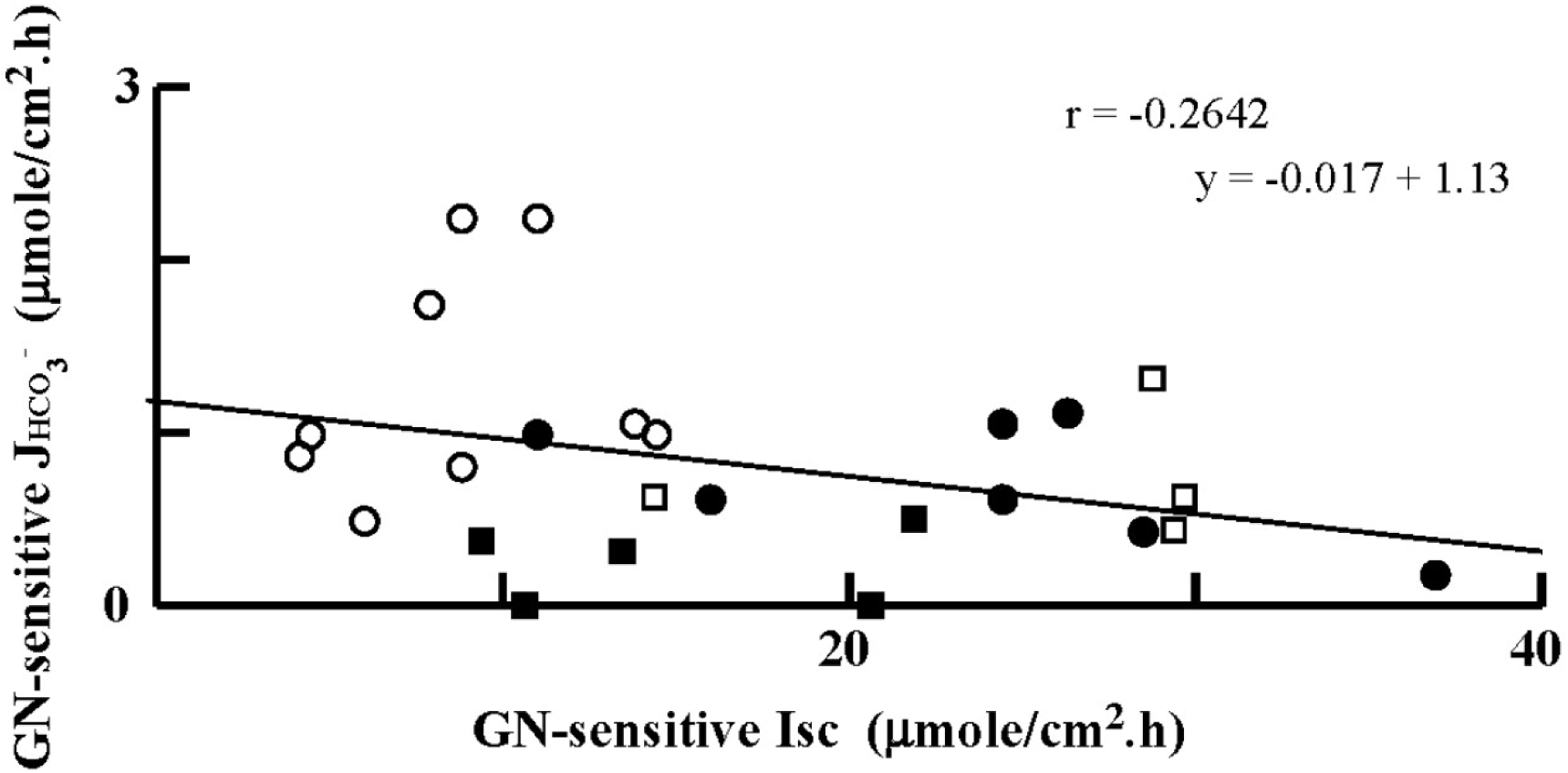
Correlation between guanylin (GN)-induced HCO_3_^-^ secretion and short-circuit current (Isc) inhibition. Data are obtained from the first (open circle), second (closed circle), third (open square) segments of middle intestine, and from the posterior intestine (closed square). Regression line and its equation are shown on the panel.

### Comparison of GN and bumetanide effects on HCO_3_^-^ secretion

As GN profoundly inhibited NKCC2 in SW eel intestine (Ando et al., 2014), effects of GN and bumetanide were compared on the HCO_3_^-^ secretion by the early part of the middle intestine where the GN effect was greatest. When applied to the mucosal side, both GN and bumetanide decreased PD and increased HCO_3_^-^ secretion to the same degree, and these effects were not additive after consecutive administration (Fig. 3). Pretreatment with bumetanide abolished the GN effect, and vice versa (Fig. 3C), indicating a common target of these drugs.

**Fig. 3.**
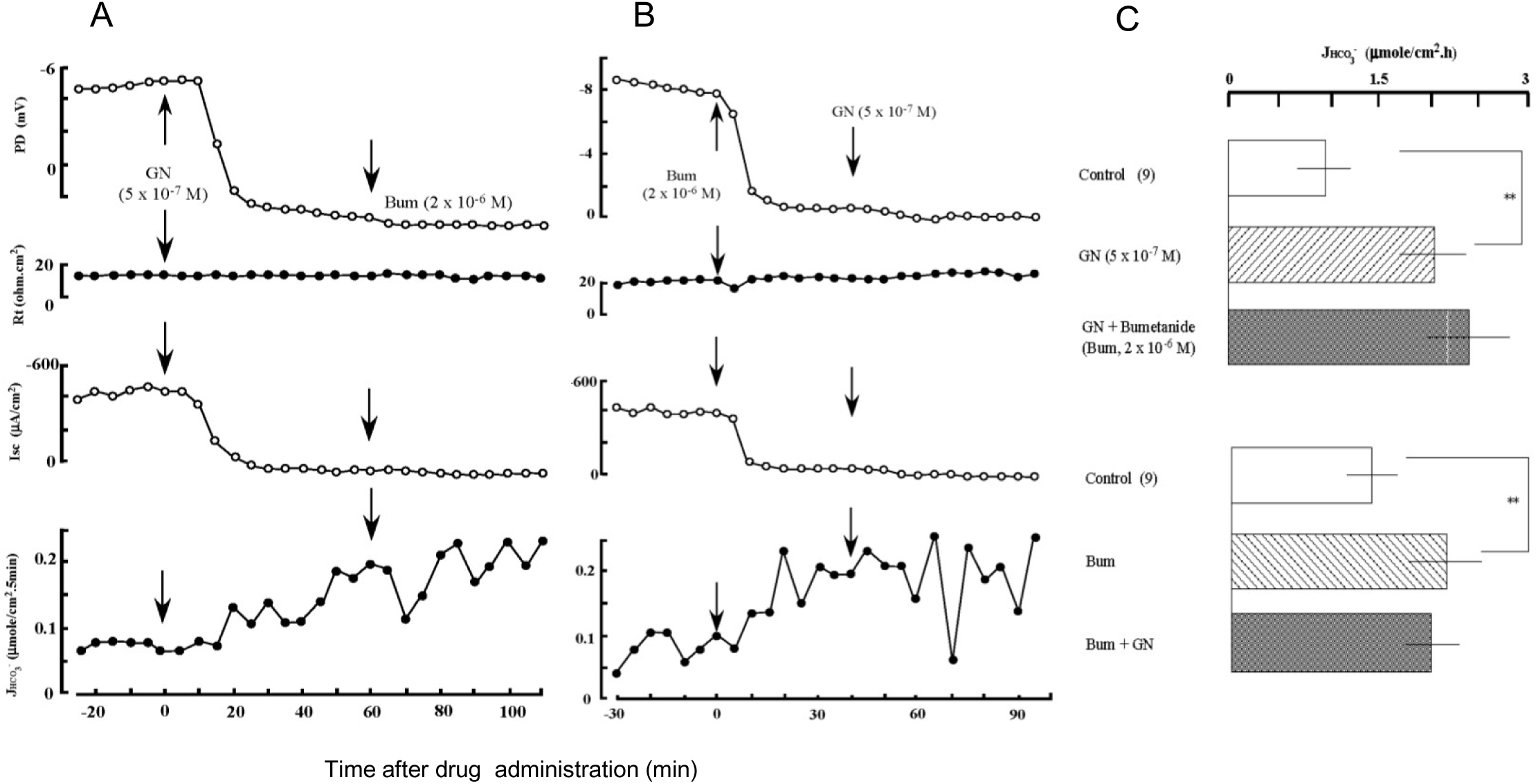
(A) An example of the effects of successive application of guanylin (GN) and bumetanide to the mucosal fluid on the transepithelial potential difference (PD), tissue resistance (Rt), short-circuit current (Isc), and HCO_3_^-^ secretion rate JHCO_3_^-^ in the middle intestine of SW eels. (B) An example of the effects of successive application of bumetanide and GN to the mucosal side on the electric parameters. (C) Summary of the successive application of GN and bumetanide, or vice versa, on HCO_3_^-^ secretion. Values are means ± SEM (n=9). **p<0.01 compared with the value before treatment.

Similar time courses were observed for GN and bumetanide actions, supporting the common target of these drugs (Table 1). Latent periods for the GN effect on PD were longer than those for bumetanide. However, the time from the PD effect to the HCO_3_^-^ secretion were similar between GN and bumetanide.

**Table 1.**
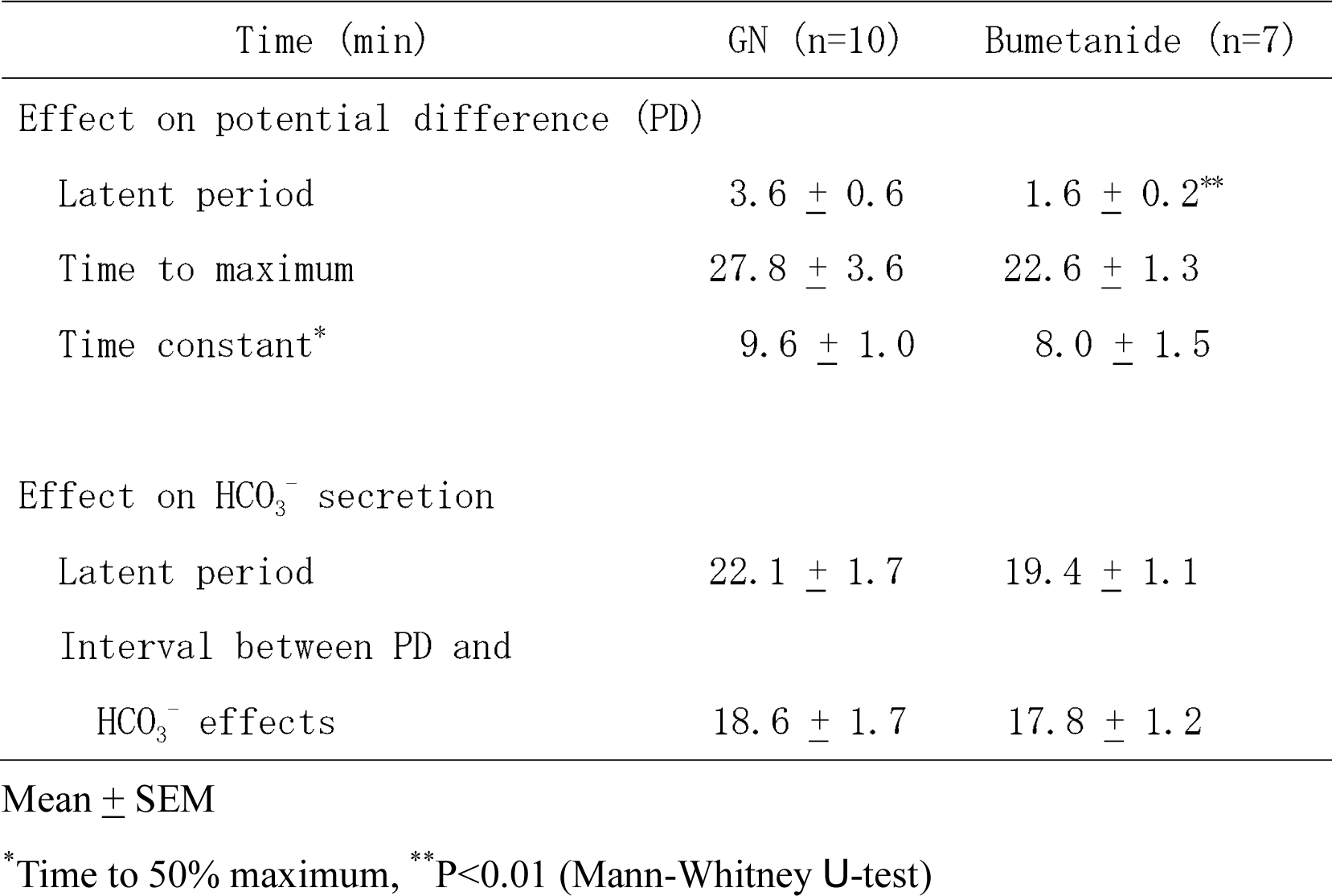
Time-course effects of guanylin (GN) and bumetanide on electrophysiological and transport parameters.

| Time (min) | GN (n=10) | Bumetanide (n=7) |
| --- | --- | --- |
| Effect on potential difference (PD) |  |  |
| Latent period | 3.6 ± 0.6 | 1.6 ± 0.2** |
| Time to maximum | 27.8 ± 3.6 | 22.6 ± 1.3 |
| Time constant* | 9.6 ± 1.0 | 8.0 ± 1.5 |
| Effect on HCO<sub>3</sub><sup>-</sup> secretion |  |  |
| Latent period | 22.1 ± 1.7 | 19.4 ± 1.1 |
| Interval between PD and HCO<sub>3</sub><sup>-</sup> effects | 18.6 ± 1.7 | 17.8 ± 1.2 |
| Mean ± SEM | | |

\*Time to 50% maximum, \*\*P<0.01 (Mann-Whitney *U*-test)

### Effects of inhibitors on GN-induced HCO_3_^-^ secretion

As the effect of DNDS was more stable and consistent than DIDS in the eel intestine, we report here only the results of DNDS treatment. Mucosal application of DNDS failed to change HCO_3_^-^ secretion, and subsequent GN application consistently increased it (Fig. 4A). The mucosal DNDS also failed to inhibit the GN-induced HCO_3_^-^ secretion (Fig. 4B). Another stilbene inhibitor, DIDS, showed similar effects (data not shown). The inhibition of apical anion channels by DPC did not inhibit the GN action when both were given to the mucosal side (Fig. 4C). However, DNDS applied to the serosal side decreased the GN-induced HCO_3_^-^ secretion gradually to a significant level (*P*<0.001) (Fig. 4D).

**Fig. 4.**
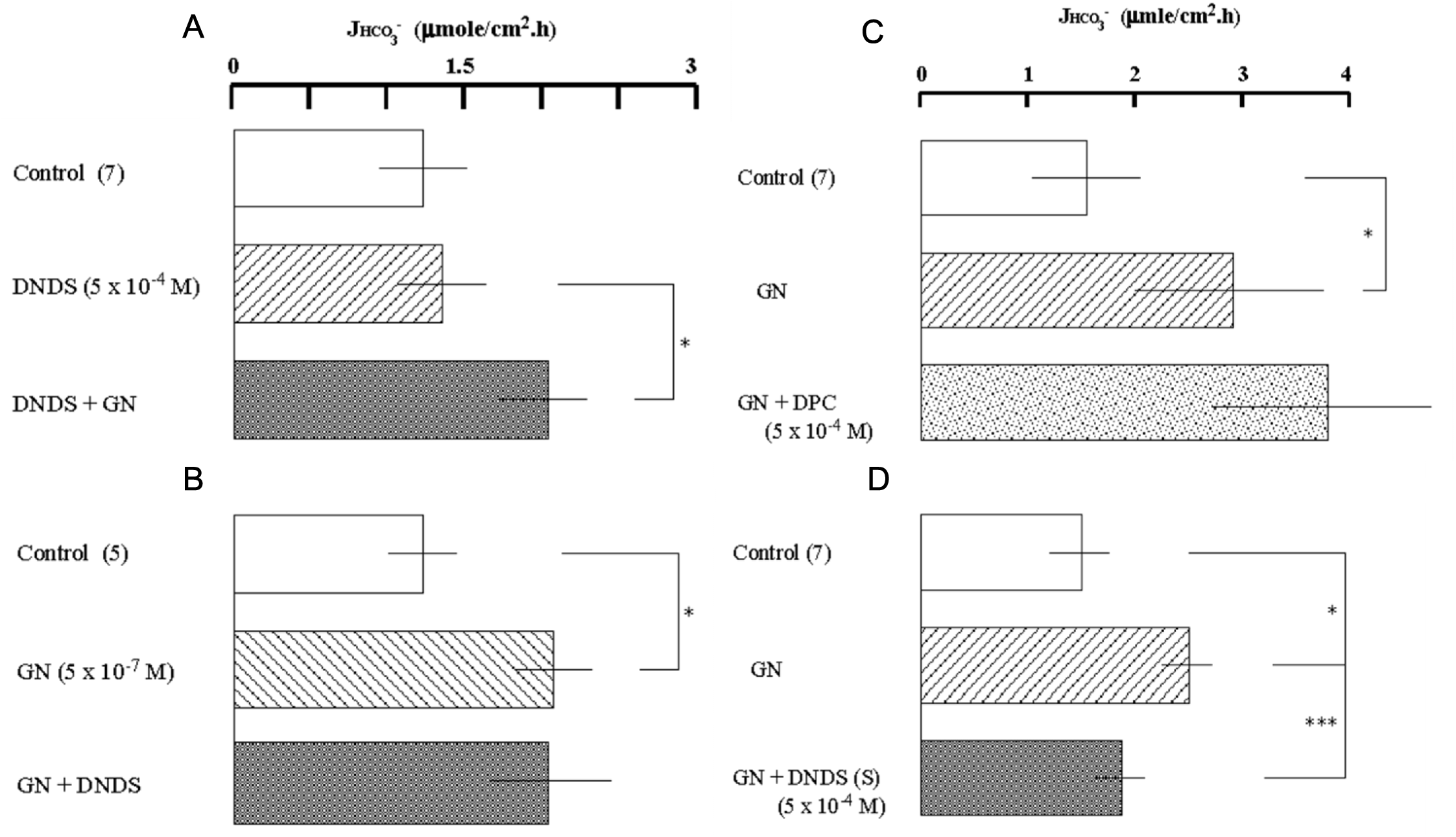
Effects of transporter inhibitors on the GN-induced HCO_3_^-^ secretion in the middle intestine of SW eels. DNDS was applied to the mucosal side before (A) or after GN (B). (C) DPC was applied to the mucosal side after GN. (D) DNDS was applied to the serosal side after GN. Values are means ± SEM. Numbers of experiments are in parenthesis. *p<0.05, ***p<0.001 compared with controls. For abbreviation of inhibitors, see footnotes.

### Changes in candidate gene expression after SW transfer

#### Transcriptomic analyses

RNA-seq analyses showed that the intestine expressed some genes of Slc4 family and Slc26 family of AE that are potentially transporters of HCO_3_^-^ from serosa to mucosa (Supplementary Table 2). Some of them are upregulated after acclimation in SW for 7 d. However, because the known paralogous isoforms (e.g. *pat1a, b*, and *c*) reported in eels could not be distinguished in transcriptome analysis, we used real-time PCR to quantify the expression of each isoform to assess their potential involvement in HCO_3_^-^ secretion.

#### Real-time PCR analyses

Gene-specific primers for each paralog were designed for qPCR based on the mRNA sequence deduced from the genome database of Japanese eel (Henkel et al., 2012) (Supplementary Table 1). For instance, two alternative transcripts were identified for Slc26a3 (DRA) and *draa* expression was more abundant in the posterior region, while *drab* expression was more abundant in the anterior region (Fig. 5). Moreover, *draa* was upregulated in the posterior intestine, while *drab* was upregulated during the course of SW acclimation in all intestinal regions, particularly in the anterior intestine. Three paralogous *pat1* were expressed in the eel intestine, and *pat1a* and *pat1b* were major transporters as suggested by their expression levels (Fig. 5). The expression of *pat1a* was greatest in the posterior intestine, while *pat1b* was expressed most abundantly in the anterior intestine. Further, the *pat1a* expression increased gradually and reached maximum after 3 d in SW, while the expression decreased for *pat1b* after SW transfer. The expression of *pat1c* increased after 3 d in SW, although the expression level was low (Fig. 5).

**Fig. 5.**
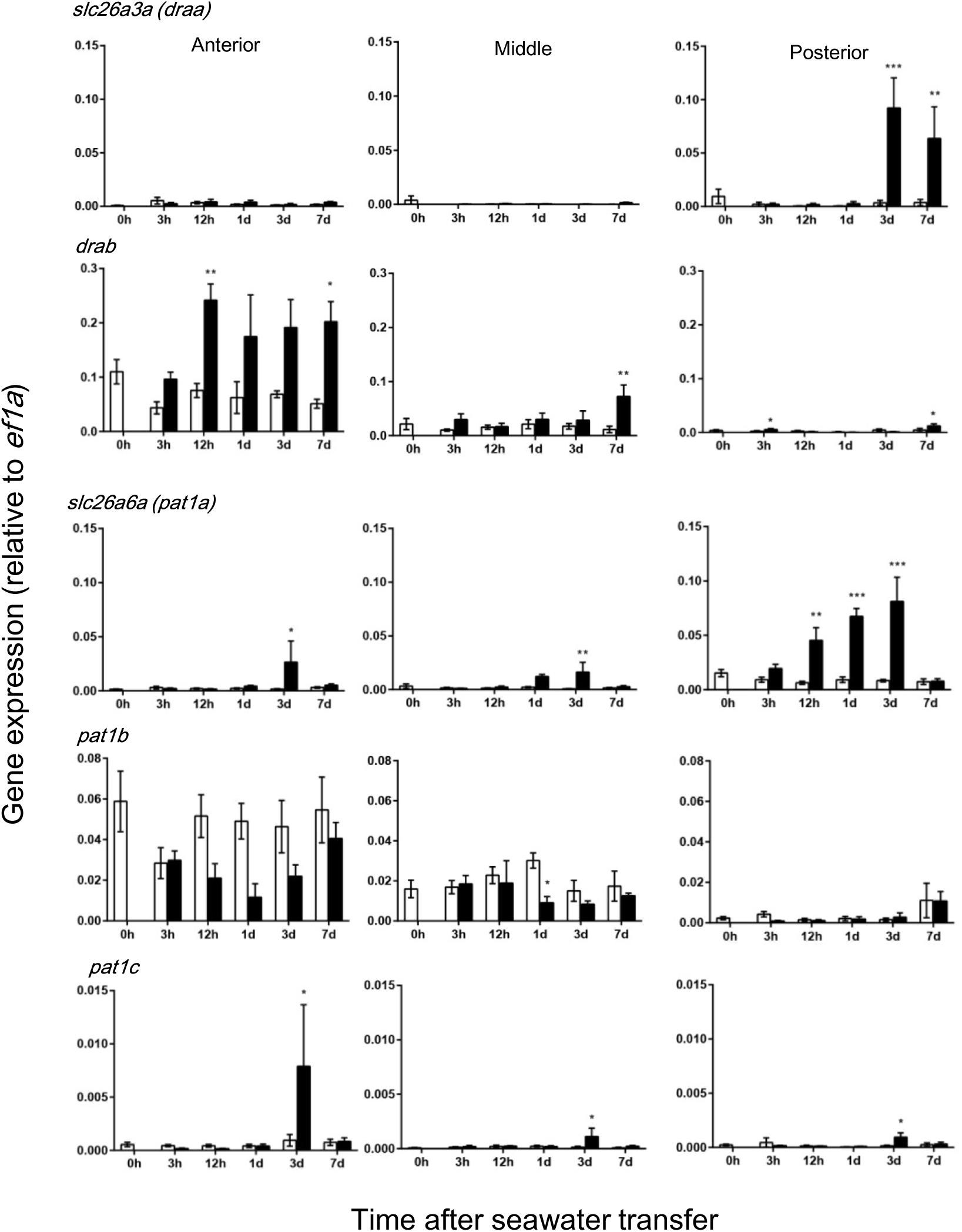
Time-course changes in gene expression of the Slc26 family of anion exchangers in different segments of eel intestine after transfer of eels from fresh water to seawater (black column) or to fresh water (controls, white column). These transporters are located on the apical membrane of epithelial cells and likely to be involved in HCO_3_^-^ secretion. Values are means ± SEM. *p<0.05, **p<0.01, ***p<0.001 compared with fresh water controls. For abbreviations, see footnotes.

Concerning basolateral HCO_3_^-^ transporters, the expression of *sat1* was greater posteriorly along the intestine, and its expression level increased 3 d after SW transfer in all segments (Fig. 6). The expression of *slc4a2a* and *b* (*ae2a, b*) increased at some time points after SW transfer, but their expression levels were very low in all segments (Fig. 6). The expression of *slc4a1* (*ae1*) was even lower than that of *ae2*s. Three *nbc* genes were expressed in the intestine, of which *nbce1a* was most abundant followed by *nbcn1* and *nbcn2a* (Fig. 7). The expression of *nbce1a* decreased along the anterior-posterior axis, while those of *nbcn1* and *nbcn2a* increased posteriorly. The expression of the *nbc* genes increased during the course of SW after acclimation in all intestinal segments (Fig. 7).

**Fig. 6.**
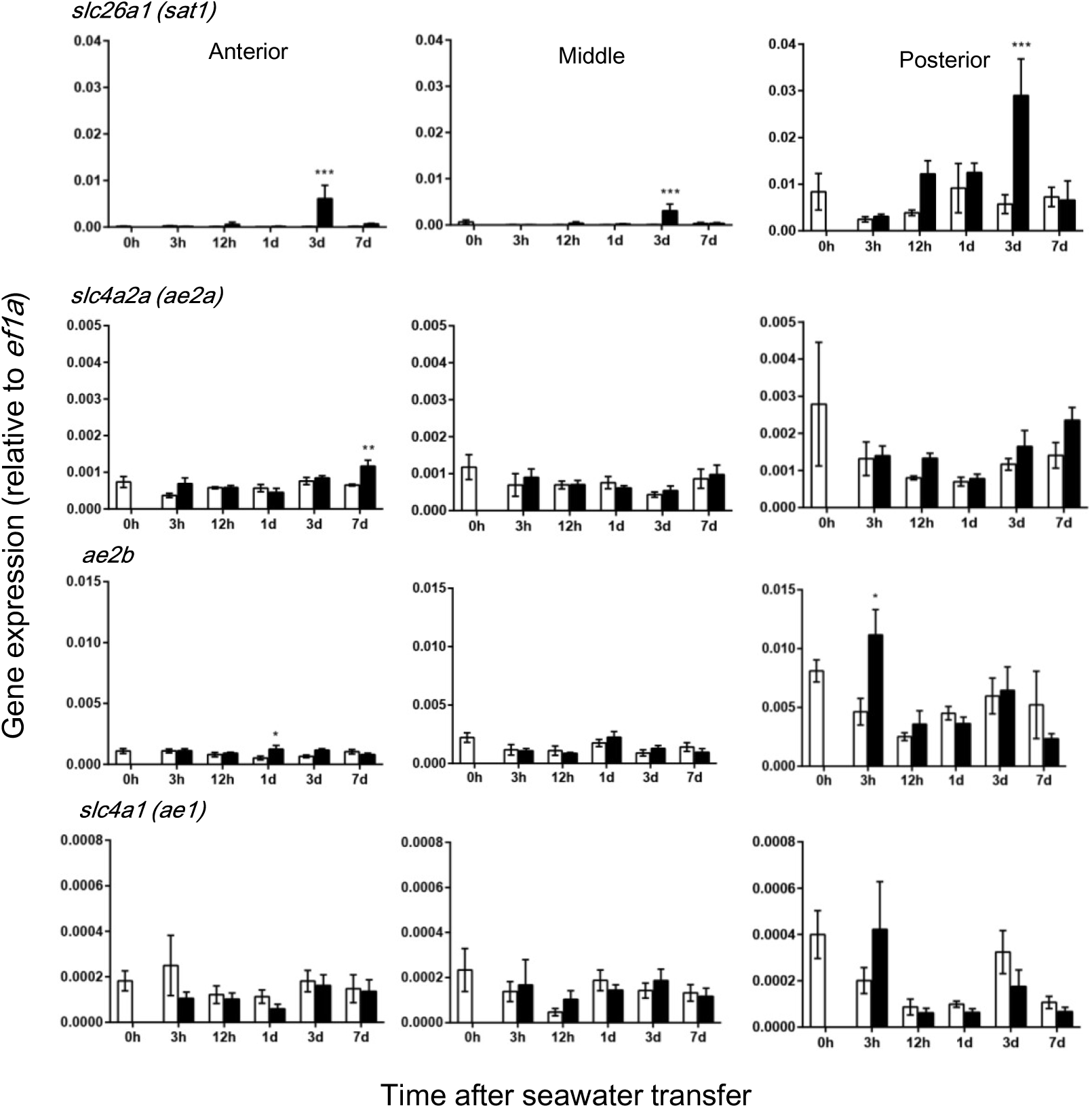
Time-course changes in gene expression of anion exchangers in different segments of eel intestine after transfer of eels from fresh water to seawater (black column) or to fresh water (controls, white column). These transporters appear to be located on the basolateral membrane and involved in HCO_3_^-^ uptake. Values are means ± SEM. *p<0.05, **p<0.01, ***p<0.001 compared with fresh water controls. For abbreviations, see footnotes.

**Fig. 7.**
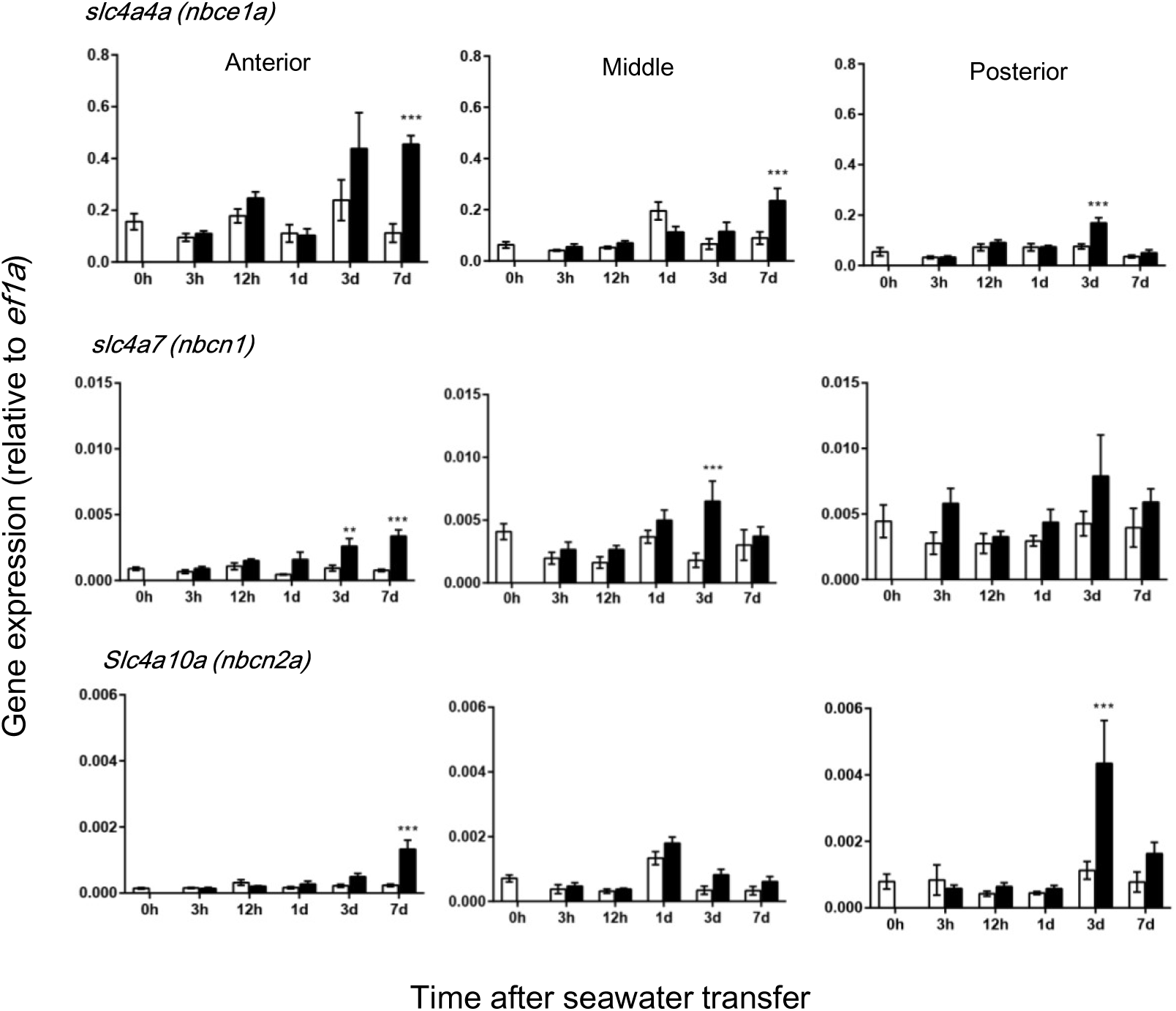
Time-course changes in gene expression of Na^+^- HCO_3_^-^ cotransporters in different segments of eel intestine after transfer of eels from fresh water to seawater (black column) or to fresh water (controls, white column). These transporters are located on the basolateral membrane and likely to be involved in HCO_3_^-^ uptake. Values are means ± SEM. *p<0.05, **p<0.01, ***p<0.001 compared with controls. For abbreviations, see footnotes.

## Discussion

The intestine is an important osmoregulatory organ for adaptation to the hyperosmotic SW environment, where water is absorbed from imbibed SW to compensate for osmotic water loss (Grosell, 2011). To achieve efficient water absorption, teleost intestine secretes HCO_3_^-^ into the lumen to remove concentrated divalent ions (Mg^2+^ and Ca^2+^) in SW as carbonate precipitates and to decrease luminal fluid osmolality (Grosell et al., 2005; Wilson et al., 2009). Hormones such as growth hormone/IGF-I and cortisol, which re-organize osmoregulatory organs to a SW-type, have long been implicated in SW adaptation (McCormick, 2001; Takei and McCormick, 2013), but investigations of the short-acting, oligopeptide hormones are still new. One of the promising candidates for SW-adapting hormone is the GN family of peptides as they are the only oligopeptide hormones thus far identified that are upregulated at both ligand and receptor levels after transfer of eels to SW (Comrie et al., 2001a, b; Yuge et al., 2003; 2006). Thus, it is quite intriguing to examine how GN regulates HCO_3_^-^ secretion through its action on the transport proteins.

### Molecular mechanisms for transcellular HCO_3_^-^ secretion

The transporters involved in HCO_3_^-^ secretion have been pursued in several teleost species. On the apical membrane of intestinal epithelial cells, Pat-1 may play an important role in HCO_3_^-^ secretion as its stoichiometry is *n*HCO_3_^-^/1Cl^-^ as shown in the euryhaline pufferfish, *Takifugu obscurus* (Kato et al., 2009), although there are conflicting reports on the stoichiometry of Pat-1 in mammals (see Alper and Sharma, 2013). In fact, *pat1* is profoundly upregulated after transfer of this euryhaline pufferfish from FW to SW (Kurita et al., 2008) and after transfer of toadfish (Ruhr et al., 2016) and sea bream (Gregŏrio et al., 2013) from SW to concentrated SW. Consistently, mucosal DIDS, but not serosal DIDS, inhibited HCO_3_^-^ secretion in the sanddab, *Citharichthys sordidus* (Grosell et al., 2001). In eels and other teleosts, however, serosal application of DIDS/DNDS, but not its mucosal application, was effective for the inhibition as confirmed in this study (Dixon and Loretz, 1986; Ando and Subramanyam, 1990; Faggio et al., 2011). Further, HCO_3_^-^ concentration of the serosal fluid greatly influences HCO_3_^-^ secretion in these teleosts. The responsible transporter for HCO_3_^-^ uptake on the serosal side has been suggested as NBCe1 in a few species of teleost because of upregulation in hyperosmotic environment (Kurita et al., 2008; Taylor et al., 2010; Grosell, 2011; Gregŏrio et al., 2013). Thus, transcellular HCO_3_^-^ transport could be conducted by apical Pat-1 and basolateral NBCe1 in some teleost species. The cytosolic HCO_3_^-^ may also be provided by hydration of CO_2_ catalyzed by cytosolic carbonic anhydrase (Grosell et al., 2009a; Gregŏrio et al., 2013).

By RNA-seq and subsequent qPCR, we found that three *pat1* paralogs were expressed in the eel intestine and *pat1a* and *pat1c* were upregulated after SW transfer in different parts of the intestine, suggesting their potential roles in HCO_3_^-^ secretion. However, the expression of two *dra* paralogs was higher than that of *pat1*, although the reverse is true for the sea bream (Gregŏrio et al., 2013). In particular, the *drab* expression was greatest in the anterior intestine and most profoundly upregulated after SW transfer, which coincides with the observation that white precipitates are observed in the lumen of anterior intestine within a day after transfer of eels to SW. In the posterior intestine, *draa* may take over the role of *drab* for HCO_3_^-^ secretion and Cl^-^ absorption as suggested by profound upregulation of the gene at this segment. Thus not only Pat-1, but also DRA is likely involved in the HCO_3_^-^ secretion in different segments of SW eel intestine. However, mucosal DNDS/DIDS failed to decrease HCO_3_^-^ secretion in this study, which suggests that eel Pat-1 and DRA are not involved in HCO_3_^-^ secretion in SW eel intestine, or these AEs of eels are insensitive to these stilbene inhibitors. It is important to note that DNDS/DIDS effectively blocked Pat-1 in mammals (Stewart et al., 2009) and teleosts (Boyle et al., 2015), but DIDS was ineffective to block DRA in the rat (Barmeyer et al., 2007) and mouse (Whittamore and Hatch, 2017). There are no studies on DIDS/DNDS sensitivity to teleost DRA, but it is possible that they are ineffective to block eel DRAs.

We identified three NBC genes in the eel intestine as basolateral transporters, *nbce1a, nbcn1* and *nbcn2a*, and they are upregulated after SW transfer. Thus, HCO_3_^-^ may be taken up from the serosal fluid by electrogenic NBCela and electroneutral NBCn1 and NBCn2a. Interestingly, stoichiometry of the human NBCe1 is 1Na^+^/2HCO_3_^-^; however, external Na^+^ concentration affects the stoichiometry in the NBCe1 of euryhaline pufferfish and the ratio reached 1Na^+^/4HCO_3_^-^ at 120 mM Na^+^ (Chang et al., 2012). Another candidate for HCO_3_^-^ uptake is Sat-1 whose gene was upregulated after SW transfer in the eel intestine. We also identified AE1 and AE2 that are usually located on the basolateral membrane of transport epithelia (Romero et al., 2013), but their expression levels were very low in the SW eel intestine.

### Mechanisms of the GN action on HCO_3_^-^ secretion

The rate of HCO_3_^-^ secretion in the SW eel intestine observed in this study seems to be higher than those of other marine fishes thus far reported (Table 2). This may be due to lower pH (7.4) used in the present study, as a linear relationship was reported between HCO_3_^-^ secretion rate and pH in the flounder intestine (Wilson and Grosell, 2003). Another possibility is higher concentration of serosal HCO_3_^-^ (24.9 mM) used in this study compared with other studies, since HCO_3_^-^ secretion across the toadfish anterior intestine is dependent on the serosal HCO_3_^-^ concentration (Taylor et al., 2010).

**Table 2.**
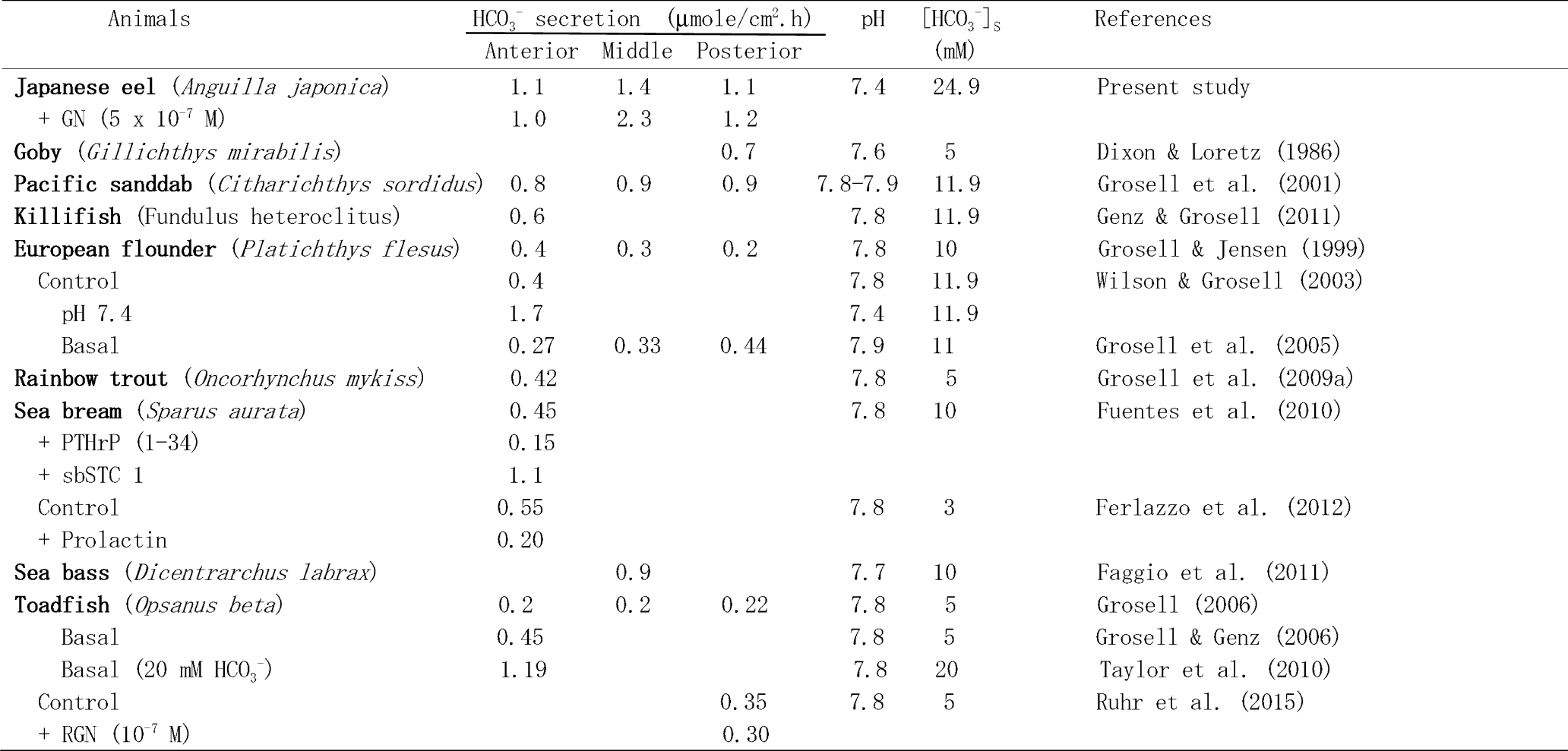
Rates of HCO_3_^-^ secretion in various segments of the intestine isolated from marine teleosts

#### Effects on the apical transporters

The present study showed that GN activates HCO_3_^-^ secretion after a latent period of about 20 min. The slow enhancement indicates that GN may not act directly on the HCO_3_^-^ transporters on the apical membrane of epithelial cells but through its action on NKCC2 inhibition (Ando et al., 2014). Consistently, the effects of GN and bumetanide were similar in both magnitude and time course, and they were not additive. It seems that the NKCC2 inhibition decreases intracellular Cl^-^ concentration, which then activates DRA/Pat-1 on the apical membrane to take up Cl^-^ in exchange of HCO_3_^-^. The upregulation of two *dra* and three *pat1* supports this hypothesis. If eel Pat-1s are electrogenic as discussed above, the serosa-negative PD should decrease after GN action. However, there was no correlation between changes in HCO_3_^-^ secretion and PD after GN, suggesting that the induced HCO_3_^-^ secretion is electroneutral. As DRA is electroneutral in mammals (see Alper and Sharma, 2013), it is possible that electroneutral and DNDS-insensitive DRAs are involved in GN-induced HCO_3_^-^ secretion in the eel (Fig. 8).

**Fig. 8.**
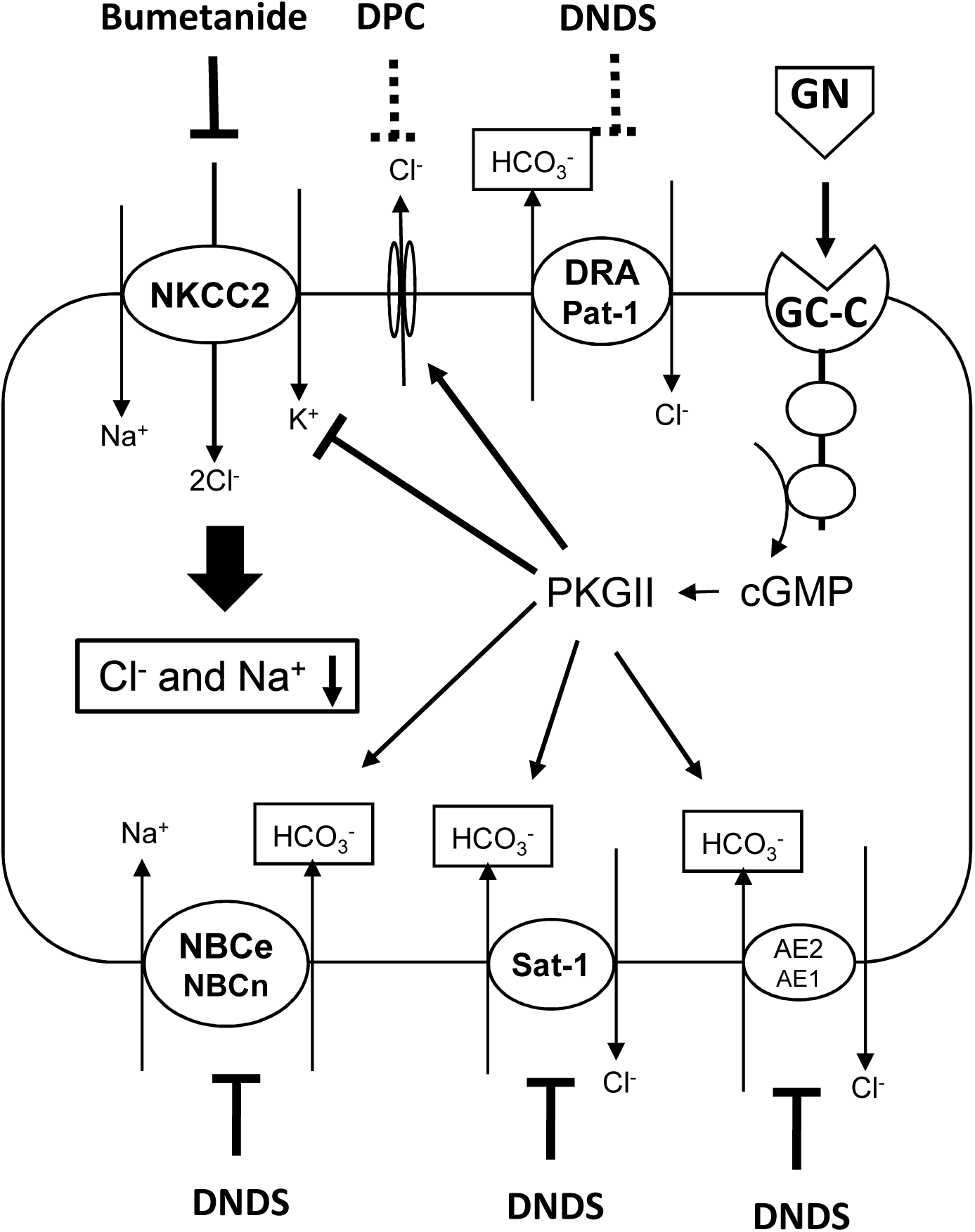
A possible mechanism of GN action on transepithalial HCO_3_^-^ secretion in the SW eel intestine. Thick lines from phosphokinase G II (PKGII) show established actions of GN on NKCC2 (inhibition) and on Cl^-^ channel (stimulation), and thin lines show deduced action on basolateral transporters (stimulation). The effect of inhibitors were effective (solid line) or ineffective (dotted line). It is possible that stimulation of Cl^-^ channel is mediated by cAMP (see text). The GN actions on NKCC2 and Cl^-^ channel decrease cytosolic Cl^-^ and Na^+^, which may stimulate apical DRA and Pat-1 and basolateral NBCs indirectly. For details, see text. Abbreviations are in footnotes.

As a second candidate for the route of HCO_3_^-^ secretion, we considered Cl^-^ channels such as CFTR, since GN stimulates apical Cl^-^ channel in the SW eel intestine (Ando and Takei, 2015) and HCO_3_^-^ is known to pass through the Cl^-^ channels (Rubenstein, 2018). Two CFTR genes exist in the eel, but the expression of the *cftr*s was low and decreased after SW transfer in the eel intestine (Ando et al. 2014; Wong et al., 2016). Considerable expression of *cftr* was reported in some teleost species (Marshall and Singer, 2002; Gregŏrio et al., 2013), and it was recently reported that eel RGN induced CFTR insertion into the apical membrane of toadfish enterocytes (Ruhr et al., 2018). However, mucosal DPC, which effectively abolished GN-induced Cl^-^ secretion in the eel intestine (Ando and Takei, 2015), failed to inhibit GN-induced HCO_3_^-^ secretion in the present study. Thus, it is unlikely that GN directly stimulates Cl^-^ channel including CFTR for HCO_3_^-^ secretion in the SW eel intestine. The slow electroneutral effect of GN on HCO_3_^-^ secretion also excludes the direct involvement of Cl^-^ channel in the GN effect. The remaining possibility is that GN further decreased cytosolic Cl^-^ through Cl^-^ channel, and the decreased cytosolic Cl^-^ activates DNDS-insensitive DRA (Fig. 8). This route may be active in SW eel intestine *in vivo*, where Cl^-^ concentration is very low and the function of DPC-sensitive Cl^-^ channel is evident (Ando and Takei, 2015).

In the intestine of mammals, GN enhances HCO_3_^-^ secretion through its action on CFTR, but the effect is much faster than that observed in the eel intestine (Guba et al., 1996; Sellers et al., 2008). The GN effect was diminished in CFTR-deficient mice or after blocking the apical Cl^-^ channels by inhibitors (Seidler et al, 1997; Zuang et al., 2007). The plausible model for GN action is that cGMP produced after binding to GC-C inhibits phosphodiesterase, resulting in cAMP accumulation and activation of CFTR for HCO_3_^-^ secretion (Sindic and Schlatter, 2006). However, as GN enhances intestinal HCO_3_^-^ secretion in the CFTR-deficient mice (Sellers et al., 2008), GN also acts on the HCO_3_^-^ transport system other than CFTR (Ko et al., 2002).

#### Effects on the basolateral transporters

In contrast to the mucosal application, serosal DNDS inhibited GN-stimulated HCO_3_^-^ secretion. This indicates that HCO_3_^-^ uptake across the basolateral membrane is a limiting step for the transcellular HCO_3_^-^ flux from serosa to mucosa. Of the HCO_3_^-^ secreted into the lumen, 80% was suggested to be taken up from the serosal fluid and the rest was produced within the cell from CO_2_ by carbonic anhydrase in the enterocyte of sea bass (Faggio et al., 2011). NBCe1 has been suggested to be responsible for HCO_3_^-^ uptake in the pufferfish (Kurita et al., 2008), toadfish (Taylor et al., 2010) and sea bream (Gregŏrio et al., 2013). Six possible HCO_3_^-^ transporters, NBCela, NBCn1 and NBCn1a, Sat-1, AE1 and AE2, appear to exist on the basolateral membrane of eel enterocytes and three *nbc* genes and *sat1* were upregulated after SW transfer. Thus it is possible that GN stimulates these basolateral transporters (Fig. 8). The uptake of HCO_3_^-^ across the basolateral membrane may explain the slow process of GN-induced HCO_3_^-^ secretion. However, it is also likely that the decrease in intracellular Na^+^ after NKCC2 inhibition stimulates Na^+^ and HCO_3_^-^ cotransport by the NBCs, which then promote apical DRA for HCO_3_^-^ secretion (Fig. 8). The intestinal HCO_3_^-^ secretion is inhibited by prolactin in the seabream, which was mediated through the downregulation of *nbce1* (Ferlazzo et al., 2012), supporting the rate-limiting role of basolateral NBCe1 in HCO_3_^-^ secretion in this species.

In the toadfish intestine, RGN contrarily inhibited HCO_3_^-^ secretion, but it increased the secretion slightly in fish acclimated in concentrated SW (Ruhr et al., 2015). The weak effect of RGN may be due to the low HCO_3_^-^ concentration of serosal fluid in their study and greater dependence on the cytosolic HCO_3_^-^ produced by carbonic anhydrase in this species (Taylor et al., 2010). In the eel, GN-induced HCO_3_^-^ secretion depends deeply on the HCO_3_^-^ uptake from the serosal fluid. Judging from the greater dependence of RGN effect on Pat-1 and CFTR in the toadfish, the molecular mechanisms for HCO_3_^-^ secretion in the intestine and its regulation by hormones are highly diverse among teleost species as observed in other osmoregulatory organs (Takei et al., 2014). Concerning other hormones, stanniocalcin stimulated HCO_3_^-^ secretion in the sea bream intestine and parathyroid hormone-related peptide inhibited it (Fuentes et al., 2010, Table 2).

## Acknowledgements

The authors thank Drs. Haruka Ozaki, Wataru Iwasaki and Yuzuru Suzuki of the University of Tokyo for transcriptomic analysis, and Dr. Christopher A. Loretz of State University of New York at Buffalo for his valuable comments on the manuscript and polishing English. This work was supported by Grant-in-Aid for Scientific Research (A) from the Japan Society for the Promotion of Science (23247010) and by Grant-in-Aid for Scientific Research on Innovation Areas “Genome Science” from Ministry of Education, Culture, Sports, Science and Technology of Japan (221S0002) to Y.T.

## Abbreviations

AE: anion exchanger;
CA: carbonic anhydrase;
CFTR: cystic fibrosis transmembrane conductance regulator;
DIDS: 4,4’-diisothiocyano-2,2’-disulfonic acid;
DMSO: dimethyl sulfoxide;
DNDS: 4,4’-dinitrostilbene-2,2’-disulfonic acid;
DPC: diphenylamine-2-carboxylic acid
DRA: down-regulated in adenoma;
FW: fresh water;
GN: guanylin;
Isc: short-circuit current;
NBC: Na^+^- HCO_3_^-^ cotransporter;
NCC: Na^+^-Cl^-^ cotransporter;
NKA: Na^+^/K^+^-ATPase;
NKCC: Na^+^-K^+^-2Cl^-^ cotransporter;
Pat-1: Putative anion transporter-1;
PD: potential difference;
Sat-1: Sulfate transporter-1;
Slc: Solute Carrier;
SW: seawater

